# Decoding the sex of faces using the power, phase, and the Fourier spectrum of the time-frequency representation

**DOI:** 10.1101/2025.07.02.662280

**Authors:** Szabolcs Sáringer, András Benyhe, Péter Kaposvári

## Abstract

The EEG activity related to face processing has been studied extensively with machine learning techniques and most of these studies apply preprocessed data without further data transformation. Here we analyzed the data of two experiments and explored the potential in decoding the time-frequency representation in face processing, thus extracting not only temporal but also the frequency distribution.

In the two experiments, participants were presented with faces and through changing the presentation number per face, we manipulated their familiarity. Then we determined the time window of facial sex related cortical activities with frequently employed decoding techniques on the preprocessed data. Subsequently, we performed Fourier transformation on the data and decoded time-frequency spectrum of the amplitude, the phase and also the complex Fourier spectrum.

This analysis revealed a 500 ms long time window at the beginning of the stimulus presentation in the 2 to 17 Hz frequency range, which showed above-chance decoding accuracies in the case of more familiar faces. Less familiar faces showed similar, albeit more restricted time-frequency windows. By comparing the two experiments we also observed 350 ms long window in the low frequencies of 4-10 Hz, where familiar faces exhibited greater decoding accuracies.

This method expanded on the generally observed time-window and complemented it with a frequency distribution related to facial sex processing. The current study demonstrates that machine learning applications can be applied to higher dimensional data, like the time-frequency representation in cognitive studies.

## 1. Introduction

Faces are one of the most important visual stimuli in humans. The neural background of face processing has remained a prominent topic in visual research and has been investigated in humans and non-humans. This investigation has identified several crucial parts of this process, like the role of the ventral stream in face recognition and the fusiform face area. A new wave of research aims to identify neural patterns associated with different aspects of face recognition and face processing using machine learning. These decoding techniques, like the multivoxel/multivariate pattern analysis (MVPA) or the representational similarity analysis (RSA), are a great way to extract and identify neural patterns associated with a predefined processing mechanism. The mentioned methods include mostly fMRI and M/EEG data but can also be applied to single-unit recordings (Anzellotti et al., 2014; Cauchoix et al., 2014; Dubois et al., 2015). A great advantage of these machine learning applications is the versatility of the input data. Pattern recognition can also be successfully applied to low-dimensional in vivo electrode recordings and high-dimensional fMRI recordings. The input data’s attributes determine the output’s main feature (Carlin, 2015). For example, fMRI data can be applied to localize given processes within the brain, while EEG is a great tool to determine the temporal distribution of said processes. A further advantage of the technique is that several aspects of a stimulus can be investigated at once. By applying different facial stimuli, various features can be decoded, such as the sex, identity, familiarity, or age of the face (Ambrus et al., 2019, 2021; Dobs et al., 2019).

Substantial contributions have been made in fMRI decoding regarding face processing, especially focusing on face-sex decoding. The sex of a face can be decoded from BOLD responses; furthermore, this process-related BOLD activity can be localized to distinct regions. One study identified the occipital and temporal regions, where above-chance decoding could be carried out, while the frontotemporal area showed no sex-related activity (VanRullen & Reddy, 2019). Another study observed that the sex feature activated the core and extensive face network (Kaul et al., 2011). These results are sometimes contradictory, since several studies failed to decode face-sex from the occipital region (Kriegeskorte et al., 2007) or the inferotemporal cortex (Kanwisher, 2017), while some found that the sex-based face-categorization is localized in the fusiform face area (Contreras et al., 2013).

Using EEG data, we can also determine the sex-related time windows of the cortical patterns presenting facial stimuli. Numerous studies have investigated this phenomenon with success using ERP data. These studies agree that sex-related activities in the brain arise with a very short latency, around 100 ms after stimulus presentation (Ambrus et al., 2019, p. 201, 2021; Dobs et al., 2019). Results also show even quicker emergence (Nemrodov et al., 2016, 2018), but this must be treated with caution since many studies apply a sliding averaging window to their data (Vida et al., 2017).

Another common feature of these studies mentioned above is that they apply the decoding methods to preprocessed event-related EEG data, without further processing it. However, MVPA and RSA techniques are more than capable of handling higher-dimensional data, such as the Fourier spectra of the time-frequency representation (TFR). Previous studies have used decoding methods on different frequency bands, but these mostly included extracting power from various ranges and testing them independently (Bae & Luck, 2019).

The decoding of the TFR is mostly popular among brain computer interface studies (Alazrai et al., 2017; Li et al., 2016; Xu et al., 2014), but it is rarely applied to cognitive EEG studies. To our knowledge, one previous study applied decoding of the TFR to observe face-processing related activity using intracranial recording in humans (Tsuchiya et al., 2008). In their study, they investigated several facial features and related cortical activity, e.g., face vs non-face stimuli, fearful vs happy expressions, or gender. Their findings included significant frequency windows below 30 Hz and between 60 and 150 Hz related to face-processing and mostly expression processing. They also found that the right hemisphere and its activity showed better decoding performance compared to the left.

Our current study focused on the exploration of the potential in decoding TFR data and its possible utilization in cognitive studies. We reanalyzed previously recorded EEG data of two experiments (Saringer et al., 2025, preprint). In Experiment 1, participants were exposed to 6 face stimuli, and each was presented 100 times. In Experiment 2, the number of faces and the repetition number were inverted, so 100 faces were presented six times. This different identity/repetition ratio resulted in procedural familiarity to different degrees in the two experiments. In the first step, we analyzed the data with the MVPA technique using the ERP data and classic cluster analysis of the power spectrum. After establishing these results, we performed a Fourier analysis on the data, and MVPA was applied to TFR. During our analysis, we also examined the different features of the TFR, and decoding was carried out on the power, on the phase, and also on the whole Fourier spectra. Our results show that the sex of a face can be successfully decoded from the TFR, which leads to identifying not only face-related time but also frequency windows.

## 2. Materials and Methods

### 2.1. Participants

Altogether, 50 healthy people with normal or corrected-to-normal vision participated in the two experiments (33 females, 24.81(±3.39) years mean age (±SD)). All participants gave written informed consent. The study protocol was approved by the Hungarian National Center for Public Health and Pharmacy (NNGYK/49248-3/2024).

### 2.2. Stimuli & procedure

We selected 100 pictures from the Chicago Face Database (Ma et al., 2015), balanced between male and female faces (all white with neutral expression). The pictures were grayscale, centered around the nose, and cut to an oval shape. Stimulus presentation was carried out in Psychtoolbox (Kleiner et al., 2007), MATLAB. All stimuli were presented on a gray background alongside a fixation cross. The stimuli were visible for 500 ms, which was followed by a jittered interstimulus interval (800-1200 ms). Participants were seated approximately 50 cm from the screen and were instructed about the task. The task was to indicate the appearance of tilted faces on the screen with a button press (Fig. 1). During both experiments, 600 stimuli were presented to each participant, 10% of which were target stimuli (tilted faces). The data from target trials were later excluded from the analysis. In Experiment 1, 6 faces (3 females and 3 males) were randomly selected from the stimulus stock and presented 100 times each. In experiment 2, all 100 faces were presented to the participant 6 times. All participants were exposed to 600 stimuli (60 targets, 540 non-targets), which lasted 15 minutes.

**Figure 1.**
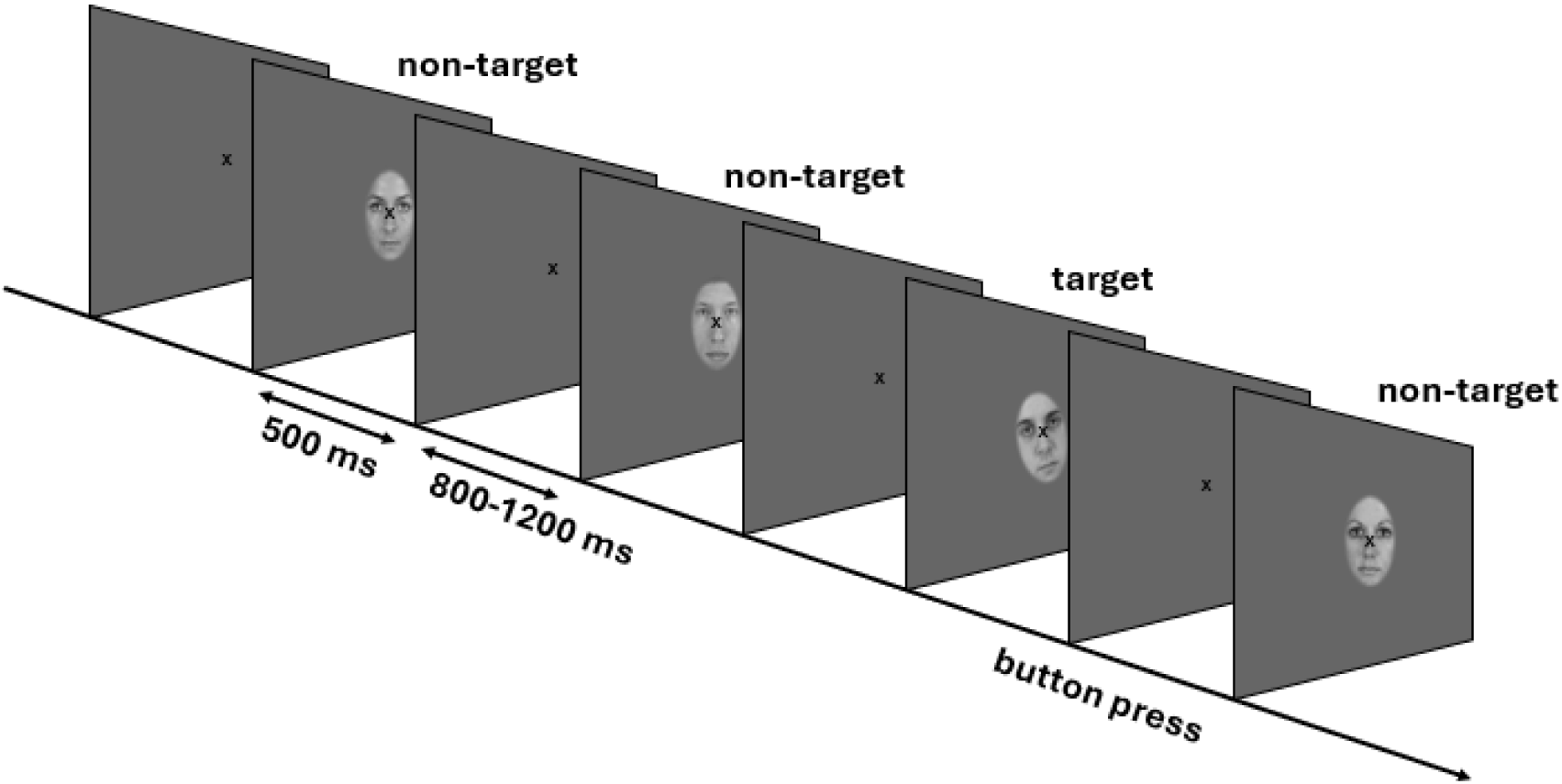
Study design: Participants were presented with 540 non-target face stimuli and 60 target stimuli (tilted faces) in a pseudorandom sequence. Each stimulus was presented for 500 ms, which was followed by a jittered interstimulus interval of 800 to 1200 ms. Participants were instructed to press a button (1 on the numeric keyboard) in case of the appearance of a tilted face as soon as they could. A button press was accepted as a hit if it happened within 1 s of the start of the stimulus presentation.

### 2.3. Data acquisition and preprocessing

During stimulus presentation, a 64-channel EEG was recorded using Biosemi Active II with a sampling rate of 512 Hz. For the purpose of later analysis, 4 EOG channels were also placed on the outer canthi and above and below the left eye.

**Figure 2.**
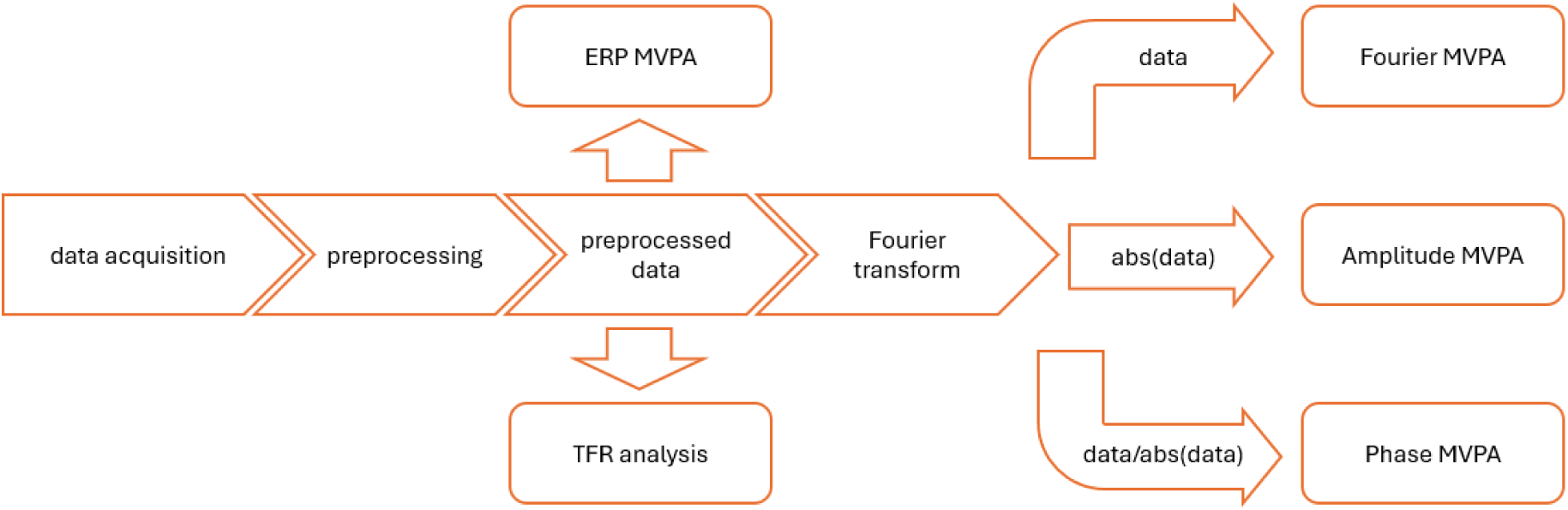
Processing pipeline. The acquired data was preprocessed, which was later analyzed using MVPA and TFR cluster analysis on the power spectra. After a Fourier transform, the time-frequency data was analyzed in three ways: the complex Fourier spectrum, amplitude by taking the absolute value of the Fourier spectra, and the phase. Phase data was acquired by dividing the Fourier spectra by their absolute value.

The preprocessing pipeline was implemented in MATLAB using the Fieldtrip toolbox (Oostenveld et al., 2010) and custom-written scripts. The data was filtered between 1 and 48 Hz. Denoising source separation was used to remove blinking and eye movements from the data. After removing artefacts, the data were epoched around the stimulus presentation: 0.5 s before the stimulus presentation and 1.2 s after. Then, the data was resampled to 100 Hz and baselined to the 0.3-0.1 s prestimulus window. As a last step, trials were removed where the data showed unusually high variance, which resulted in the exclusion of 2.4 trials per participant on average.

### 2.4. Data analysis & statistics

In the behavioral task, participants’ responses to the target stimuli were accepted if the reaction time was no longer than 1 s, measured from the beginning of the stimulus presentation. The accuracy of the detection task was used to evaluate the engagement of the participants. Subjects with an accuracy score under 0.8 were to be excluded.

The preprocessed EEG (ERP) data were first analyzed using Multivariate Pattern Analysis (MVPA) implemented in the MVPA-Light toolbox (Treder, 2020). Decoding accuracy was determined at all time points from −0.1 s to 0.8 s relative to the stimulus presentation using the Linear Discriminant Analysis (LDA) algorithm. The time window was selected based on previous publications (Ambrus et al., 2019, 2021; Dobs et al., 2019), since facial sex related cortical activities have been found between 50-800 ms after stimulus presentation, approximately. LDA was chosen over the Support Vector Machine (SVM) due to the heightened computing load of time-frequency representation, and the LDA provided shorter computational durations. For cross-validation, the Leave-One-Out Cross-Validation (LOOCV) method was used, because it provides the most comparable results between the experiments.

Classic TFR analysis was implemented in the Fieldtrip toolbox (Oostenveld et al., 2010), MATLAB. Convolution with Hanning taper was carried out on the data with a fixed window length of 0.5 s between −0.1 s and 0.8 s, in the frequency window of 1 to 45 Hz. The frequency window of interest was selected based on the previously mentioned publication, where activities were observed under 50 Hz. The value of 45 Hz was determined to avoid mains interference. The acquired power spectrum was baselined to the −0.3 to −0.1 s prestimulus window, and the power is reported in dB.

To implement TFR decoding, the preprocessed data were first Fourier transformed similarly to previously described in the classic TFR analysis. The difference is that not only the power spectra but the whole Fourier spectra were extracted as a complex number. To observe the cortical patterns solely in the amplitude, we took the absolute value of the whole Fourier spectra. To observe the phase patterns, the Fourier spectra were divided by the absolute value of the data. This allowed us to observe the phase data by normalizing the amplitude at all time points; furthermore, this way we circumvented the circular nature of the phase data. MVPA was carried out in all three approaches in the same manner: the LDA algorithm was trained with LOOCV to acquire decoding accuracy at all times and frequencies.

Statistical significance was determined in the same way in all analyses. In the TFR analysis trials with female faces were compared to trials with male faces using paired t-tests. Threshold-free cluster enhancement (TFCE) was used for correction, and significance was determined against a population created from a permutation of 10,000 iterations. In the MVPA analyses, the decoding accuracy value was compared to chance performance (0.5) at all data points; then corrected with TFCE (10,000 iterations).

## 3. Results

### 3.1. Experiment 1

All participants’ performance on the detection task was above 0.8, on average: mean (±SD) = 0.97 (±0.03) with a reaction time of: mean (±SD) = 0.50 (±0.06) s. Based on the behavioral data, participants’ engagement was evaluated as high, and no participant was excluded.

**Figure 3.**
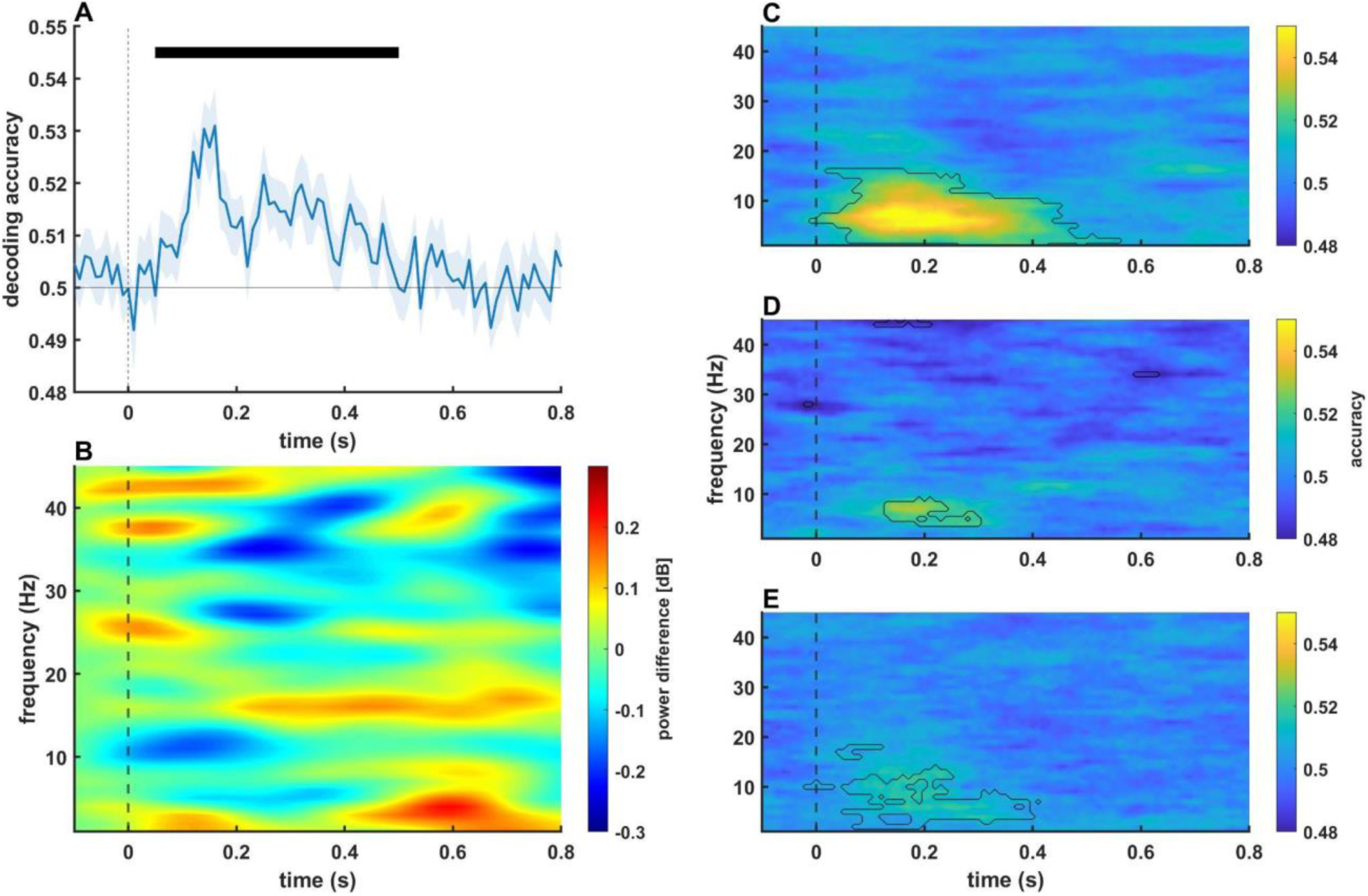
The results of Experiment 1. A: ERP decoding across time, where the blue line represents the average decoding accuracy and the shaded area marks the SEM. Significant time windows are indicated by the horizontal black bar. B: TFR analysis, where the power difference between the two conditions is visible across time and frequency. C, D, E: Decoding accuracy using the Fourier spectrum, the amplitude, and the phase, respectively, across time and frequency. Above-chance performance of the algorithm is indicated by the black contour (p<0.05)

The cluster analysis of the power spectra revealed no significant cluster in the power difference of the conditions (p>0.05). Two narrow bands in the time-frequency window between 0 and 200 ms around 10 Hz and between 200 and 700 ms around 16 Hz were present after TFCE correction, but neither cluster reached significance. In the case of the earlier window, female faces showed higher power, while the opposite can be observed in the latter cluster. The MVPA of the preprocessed EEG data showed a significant time window between 60 and 490 ms (p<0.05).

The greatest significant cluster emerged in the case of decoding the complex Fourier spectrum. This analysis resulted in the cluster expanding from 0 to 560 ms between 2 and 17 Hz (p<0.05). The other two tests revealed clusters with more limited distributions. Decoding of the amplitude data showed a cluster extending in the time frequency window of 130 to 300 ms and 4 to 10 Hz (p<0.05). The phase decoding exhibited a more diffuse distribution (0 to 400 ms, 2 to 19 Hz, p<0.05), but the data appeared to be heterogeneous with varying, low decoding accuracies.

### 3.2. Experiment 2

In Experiment 2, behavioral results were also satisfactory for all participants. Mean (±SD) detection accuracy was 0.96 (±0.03), and no participant was under 0.8. Mean reaction time (±SD) was 0.48 (±0.05).

**Figure 4.**
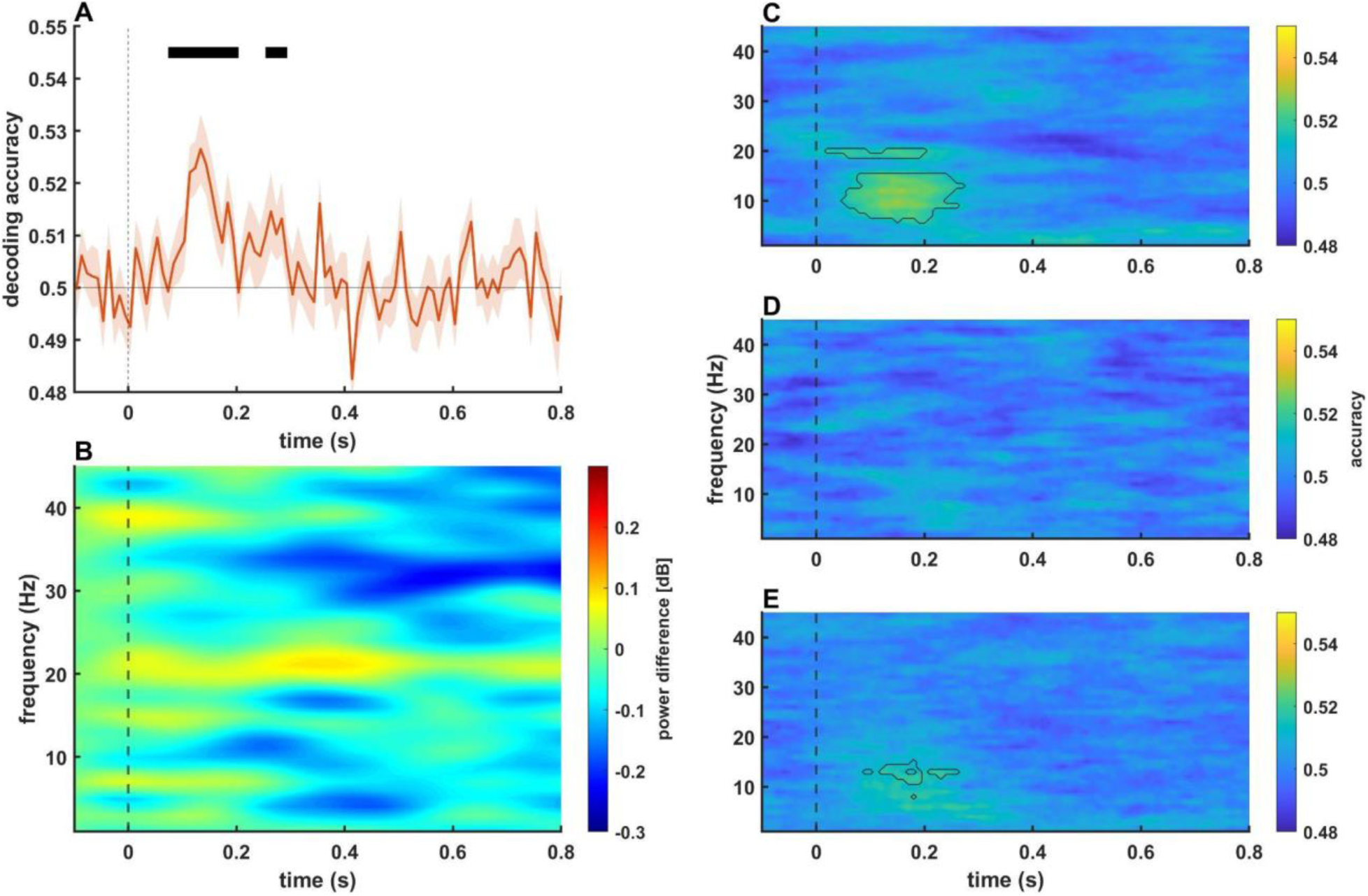
The results of Experiment 2. A: ERP decoding across time, where the orange line represents the average decoding accuracy and the shaded area marks the SEM. Significant time windows are indicated by the horizontal black bar. B: TFR analysis, where the power difference between the two conditions is visible across time and frequency. C, D, E: Decoding accuracy using the Fourier spectrum, the amplitude, and the phase, respectively, across time and frequency. Above-chance performance of the algorithm is indicated by the black contour (p<0.05)

Analysis of the power spectra did not show any significant cluster in Experiment 2, either (p>0.05). Investigating the statistical values across time and frequency showed a cluster between 500 and 700 ms at 30 and 35 Hz. This activity showed greater power in the case of the presentation of female faces. The MVPA time course of the ERP data showed comparable results to Experiment 1, with more restricted effects. Significance emerged between 80 and 280 ms, but the time window was not continuous (80 to 190 ms and 260-280 ms, p<0.05).

Alike in Experiment 1, the greatest effect emerged in the complex Fourier spectrum. Two significant clusters emerged: one spreading from 50 to 270 ms poststimulus in the frequency band of 6 to 15 Hz, and one narrow band emerging at 20 Hz spanning from 20 to 210 ms (p<0.05). In Experiment 2, the decoding of the amplitude data did not yield any significant cluster in the TFR. In contrast, the phase data decoding showed a dispersed above-chance accuracy across 90 to 260 ms in the frequency band of 10 to 15 Hz.

### 3.3. Comparison of the two experiments

To further expand on the difference between the two experiments, decoding accuracies were compared across experiments. We did not pursue the comparison of the power spectra since neither experiment showed significant effects.

**Figure 5.**
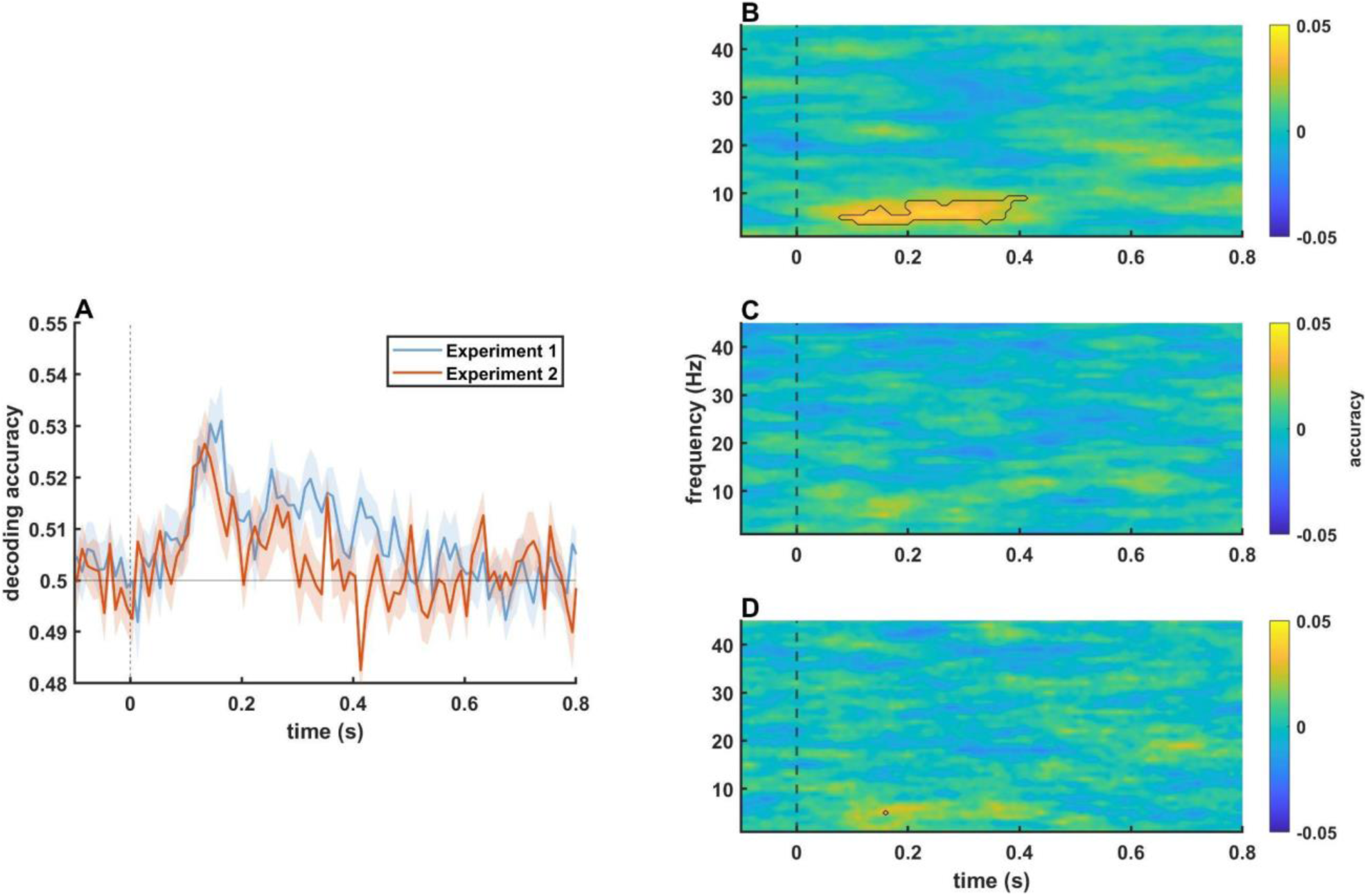
Comparison of the two experiments. A: Decoding accuracy across time in the two experiments using preprocessed EEG data. B, C, D: The difference in decoding accuracies across time and frequency between the two experiments. The black contour marks the significant cluster.

Experiment 1 yielded greater decoding accuracies and longer significant time windows, but the comparison between Experiment 1 and Experiment 2 did not show a significant difference across time. The comparative analysis resulted in a significant difference in the complex Fourier spectrum only. The significant cluster of the difference spanned from 80 to 420 ms after stimulus presentation in the frequency band of 4 to 10 Hz (p<0.05). Neither the amplitude nor the phase data comparison yielded significant differences between the two experiments.

## 4. Discussion

In the current study, we reanalyzed and compared the EEG data of two experiments. In Experiment 1, participants were exposed to six faces, each face repeated 100 times, while in Experiment 2, the ratio of the presented faces and the number of presentations per face was reversed. The goal of the reanalysis was to explore the possibility of applying machine learning to the time-frequency representation of EEG data and whether, by decoding the TFR, we could extract further information about face processing. To achieve this, we performed routinely utilized techniques, namely MVPA of the preprocessed EEG data and TFR cluster analysis of the power spectra. Additionally, we performed MVPA on the TFR data in three ways: on the complex Fourier spectra, the amplitude data, and the phase data.

In Experiment 1, where the faces were presented substantially more times and thus were more familiar, MVPA of the ERP data revealed a time window between 60 and 490 ms. Important to note here, that the length of stimulus presentation was 500 ms, and the disappearance of the stimulus could alter the cortical activity, thus impeding the facial sex related signals. The cluster analysis of the power spectra appeared to be futile, since it did not yield any time-frequency clusters when comparing the two conditions. The decoding of the TFR, however, showed promising results; the complex Fourier spectra exhibited the greatest potential, showing a cluster that arose in a nearly identical time window of 0 to 560 ms. This analysis also led us to the observation that facial sex related oscillations appeared in the 2 to 17 Hz frequency band. The amplitude and phase data also yielded significant time-frequency clusters, although more constricted. The beginning of the time-window must be treated with caution though, due to the smoothing effect of Fourier transformation, since the sliding window reduces the punctuality of the temporal resolution.

Experiment 2 showed similar results, albeit more limited in time and frequency. Classic MVPA showed a significant time window from 80 to 280 ms, and like in Experiment 1, TFR cluster analysis did not yield significant results. In this case also, the Fourier spectrum showed the greatest effect out of the three, which showed a significant time-frequency window of 20 to 270 ms poststimulus and between 6 to 20 Hz. Comparing the results of the two experiments also revealed a window only in the complex Fourier spectrum between 80 and 420 ms in the narrow band of 4 to 10 Hz, where decoding accuracy was higher in Experiment 1.

To our knowledge, decoding of TFR is mostly used in brain-computer interface studies (Alazrai et al., 2017; Bae & Luck, 2019; Xu et al., 2014), while its utilization is quite sparse in cognitive neuroscience. Examples of decoding the time-frequency spectrum can be found in the literature (Ieracitano et al., 2019; Santaji & Desai, 2020; Sreeja et al., 2017, for review see Saeidi et al., 2021), but its use has not become widespread yet, despite its high potential. EEG decoding in cognitive neuroscience is mostly used to investigate the temporal distribution of certain cortical activity. Several studies have investigated the time course of facial sex decoding, and the current results suggest an early activity where the processing of female and male faces differs. Previous studies found the first evidence around 100 ms, and it was maintained for 500 - 600 ms (Ambrus et al., 2019, Dobs et al., 2019). It is also suggested that familiarity affects the cortical processing mechanism reflected in the difference in decoding accuracy comparing familiar and unfamiliar faces (Dobs et al., 2019). Our results align well with the previous results, where a very early onset was found in both experiments. How long this activity is maintained, however, is inconclusive. In our study, faces with low familiarity elicited shorter time windows. Despite this disparity, the same temporal distribution could be decoded from the Fourier spectrum in both experiments. Furthermore, TFR decoding was also able to detect differences between the cortical activity of the two experiments, even in overlapping windows, suggesting the previously mentioned altered processing mechanism due to different levels of familiarity.

Although TFR decoding did not provide additional information about the temporal distribution of facial sex processing, the technique was able to demonstrate the frequency distribution of the observed activity. The role of the different frequency bands in face processing has been studied extensively, and most bands have been associated with different aspects of the processing mechanism (Güntekin & Başar, 2014). Changes in the delta band (1-4 Hz) have been observed in many cognitive studies, especially the increased delta power in the frontal areas due to a cognitive load (Güntekin & Başar, 2016, Harmony, 2013). Naturally, the delta band also plays a role in face processing, and various studies have found activity differences. Interestingly, the face-associated delta activity arose in the occipital region rather than in the frontal (Başar et al., 2007). These changes appeared in various aspects of face processing, such as face familiarity (Sakihara et al., 2012), Thatcherized faces (Gersenowies et al., 2010), or facial expressions (Knyazev et al., 2009; Zhang et al., 2013). We can also find evidence that the theta band (4-7 Hz) contributes to face processing, especially in face familiarity (Crespo-Garcia et al., 2012; Miyakoshi et al., 2010; Zion-Golumbic et al., 2010). This role in face familiarity aligns well with the well-known connection of the theta band and mnemonic processes (Herweg et al., 2020). The alpha band (8-12 Hz) has been mostly identified in emotion and facial expression processing (Balconi, Brambilla, et al., 2009; Balconi, Falbo, et al., 2009; Güntekin & Basar, 2007; Zion-Golumbic et al., 2010), while the beta band (13-30 Hz) has its connection with face familiarity (Özgören et al., 2005; Sakihara et al., 2012). The gamma band (>30 Hz) also appears in relation to face familiarity and expressions, but it also appears to be sensitive to face stimuli, since higher gamma powers (29-45 Hz) have been observed in faces compared to other stimuli, like inverted faces (Keil et al., 1999). These frequencies, however, do not always separate precisely. The modulation of a 5 to 20 Hz activity has been reported spanning three frequency bands, which appeared at the early stages of stimulus processing, between 50 to 300 ms after stimulus presentation (Rousselet et al., 2007). The temporal and frequency distribution of this activity aligns well with our results, suggesting an important role in face processing of this particular activity. This time-frequency window also appeared in both experiments, faces with lower and higher familiarity. A previous study has implemented an analysis similar to the one presented here, which also investigated face-processing related time-frequency components using machine learning algorithms (Tsuchiya et al., 2008). ECoG data analysis revealed that face decoding (compared to patterns) was critical in the 50 to 150 Hz range, while expressions are coded in the 60 to 150 Hz and below 30 Hz range. These activities are located in the ventral temporal cortex and tied to the 100 to 500 ms poststimulus window. They also observed the gender decoding mechanism, which was tied to the right fusiform gyri, but in the case of gender, a specific time-frequency window was not investigated. In our study, the facial sex related activity spanned a similar time window with a wide, below 20 Hz frequency range, but our study lacks any temporal distribution of this activity.

## Conclusion

In this study, we explored the possibility of using machine learning on time-frequency representation, focusing on cognitive processes, to be more precise face-processing. Through 2 experiments, we compared TFR decoding with generally used decoding applications in face processing and facial sex coding in the cortex. This analysis revealed that facial-sex-related activities arose in the first 500 ms of stimulus presentation, below 20 Hz. This method could be an interesting possibility to expand the information extraction of machine learning analysis, especially in cognitive neuroscience, where machine learning analyses are already frequently applied.

## Conflict of interest

The authors have no conflict of interest to declare.

## Data availability

Raw data and the analyzing software are available at the Open Science Framework repositorium (https://osf.io/yu4dh/).

## Acknowledgement

We would like to thank Eszter Domboróczki, our undergraduate student for her help in data acquisition and participant recruitment.

## Notes

https://osf.io/yu4dh/

### Competing Interest Statement

The authors have declared no competing interest.

